# Differed cell division angle, position of cell proliferative area, and localization of ANGUSTIFOLIA3 in lateral organs

**DOI:** 10.1101/2021.07.09.451862

**Authors:** Ayaka Kinoshita, Makiko Naito, Hirokazu Tsukaya

## Abstract

Leaf meristem is a cell proliferative zone present in the lateral organ primordia. In this study, we investigated how the proliferative zone affects the final morphology of the lateral organs. We examined how cell proliferative zones differ in the primordia of planar floral organs and polar auxin transport inhibitor (PATI)-treated leaves from normal foliage leaf primordia of *Arabidopsis thaliana* with a focus on the spatial accumulation pattern of *ANGUSTIFOLIA3* (*AN3*), a key element for leaf meristem positioning. We found that organ shape changes by PATI treatment were correlated to the angle of the cell division plane relative to the leaf primordia axis in the leaf meristem (cell division angle), but not with leaf-meristem positioning, size of the leaf meristem, or the localization pattern of AN3 protein. In contrast, different shapes between sepals and petals compared with foliage leaves were associated with both altered meristem position associated with altered *AN3* expression patterns and different distributions of cell division angles. These results suggest that lateral organ shapes are regulated via two aspects: position of meristem and cell division angles

**Summary statement:** Different lateral organs with different morphology possess different properties of meristems; cell division angles, position of cell proliferative area and AN3 localization patterns.

## Introduction

The shape of leaves plays an important role in determining their photosynthetic function. Moreover, the shape of floral organs, which are evolutionarily derived from leaves, is also important for reproductive success. Therefore, the shape of these lateral organs varies among species to maximize their survival ability in their natural habitats. To understand these variations, it is important to assess their developmental properties.

The center of morphogenesis in plants is the meristem, where active cell division occurs. The shoot apical meristem (SAM) is essential to produce new above-ground organs. In terms of lateral organs, especially for leaf primordia, the leaf meristem supplies cells to the leaf blade; thus, researchers have investigated the nature of the meristem to understand the morphogenesis of lateral organs (e.g. Esau, 1977; Donnelly et al., 1999; Kazama et al., 2010; Ichihashi et al., 2011).

In many angiosperms, such as *Arabidopsis thaliana*, the leaf meristem is at the base of each leaf (Tsukaya, 2014; 2021). Cell proliferation initially occurs throughout the leaf primordium but is restricted to its basal regions as the leaf primordium grows further (Donnelly et al., 1999; Kazama et al., 2010). The control on the restriction of the cell proliferation zone is not completely understood. However, a transcriptional coactivator called ANGUSTIFOLIA3 (AN3)/GRF-INTERACTING FACTOR1 (GIF1), which encodes a protein that is homologous to the human synovial sarcoma translocation protein (Horiguchi et al., 2005), is considered to positively control cell proliferation in leaf primordia. The spatial patterns of AN3/GIF1 (simplified to AN3, hereafter) accumulation at the base of leaf primordia match the cell proliferation zone, suggesting that AN3 may act as an important determinant in positioning the leaf meristem (Horiguchi et al., 2005; Kawade et al., 2017). Cell division angles in the *AN3*-expressing region, except for areas along the margin and vasculature, were observed to be randomized (Yin and Tsukaya, 2016), which may contribute to the two-dimensional expansion of the leaf lamina. AN3 protein moves cell-to-cell (Kawade et al., 2013), forms a gradient along the proximal-distal axis on the leaf, with the leaf base presenting the highest concentration of AN3 protein (Kawade et al., 2017) and thus is involved in the positioning of the leaf meristem. However, how *AN3* expression is restricted to the basal part of leaf primordia is still unknown (Tsukaya, 2021).

AN3 seems to be also involved in the morphogenesis of each floral organ (Lee et al., 2009, 2014). A petal in the *an3* mutant has a smaller number of cells and a narrower shape than that of the wild type (Lee et al., 2009), as seen in the foliage leaves, suggesting a common role of AN3 in leaf and petal primordia. However, the past studies indicated that the position of the meristematic activity in the petal primordia is marginal/apical in *A. thaliana* (*e.g.,* Dinneny et al., 2004), which is clearly different from the leaf primordia. Precisely said, the cell proliferation is observed in the entire primordia of the petal organ at first, and then in the distal region in later stages, differed from that of leaf primordia (Disch et al., 2006; Anastasiou et al., 2007). Thus, if AN3 has the same role in the positional determination of the meristematic zone in petal primordia, AN3 proteins are expected to accumulate apically and not basally in petal primordia; however, no previous studies have examined it. Marginal/apical positioning of the meristematic zone in petals may be an ancestral character that is directly comparable to the apical positioning of SAM (Boyce, 2007). Although the basal positioning of the leaf meristem is common in angiosperms, marginal/apical positioning in leaf primordia is also known in some ferns and gymnosperms. Therefore, a comparison of leaf primordia with floral organ primordia of the same species, with special emphasis on *AN3* expression, may contribute toward understanding the roles and evolutionary history of differences in the positioning of cell proliferation activity in the primordia of these lateral organs.

In relation to the above-mentioned positioning of the meristematic zones in lateral organs, venation pattern is also an evident difference between the leaves of angiosperms, ferns, and gymnosperms (Boyce, 2007). In the lateral organs of ferns and gymnosperms harboring meristems along the apical margin, the leaf vein shows a bifurcated pattern and is open at the end. In contrast, lobed and closed patterns are common in eudicots including *A. thaliana*; parallel patterns are common in monocots (Dengler and Kang, 2001). Based on this correlation, an interaction between spatial control of the leaf cell proliferation zone and leaf venation pattern has been suggested (Boyce, 2007).

How the spatial control of the leaf cell proliferation zone and leaf venation patterns are interconnected? In addition, are these indeed correlated? If we refer to the earlier-mentioned AN3, the cell division orientation is randomized in leaf meristem, where AN3 localized, except for the local regions along veins and leaf margin where active auxin flow is recognized (Yin and Tsukaya, 2016). This might suggest a missing link between the spatial control of the leaf cell proliferation zone and leaf venation pattern, but no prior studies have investigated this possibility.

In vein development in the leaf, biosynthesis and transportation of the plant hormone auxin (indole-3-acetic acid, IAA) plays an important role (Cheng et al., 2006). For example, *GNOM* and *PIN-FORMED* (*PIN*) genes that control polar auxin transport (PAT) (Verna et al., 2019) regulate vein formation during leaf development. Members of the PIN family have been extensively studied as major factors in PAT, the most famous being PIN1 (Okada et al., 1991). Mutations in many *PIN* genes induce defects in leaf vein patterns, with wide and bifurcated midveins and altered leaf blade shapes (Sawchuk et al., 2013). Similarly, mutants with an abnormal venation pattern mostly show altered leaf shapes in *A. thaliana* (Candela et al., 1999).

Many PAT inhibitors (PATIs) have been used to examine the role of PAT in plant organogenesis, including 2,3,5-triiodobenzoic acid (TIBA) and N-1-naphthylphthalamic acid (NPA). Through indirect evidence, they bind to the same auxin efflux carriers to inhibit their activity (Teale and Palme, 2018). When plants are treated with PATI, their leaves exhibit abnormal leaf venation patterns, including very thick midveins and marginal veins, similar to that of *pin1* mutants (Sieburth, 1999). In addition, the leaf shape becomes rounder and shorter than that of control plants. However, to date, no study has examined the effect of PATI on the cell proliferation pattern in leaf primordia.

To fully understand the morphogenesis process of lateral organs, it is necessary to investigate the role of the lateral organ meristem on the final organ shape and factors that affect the properties of meristems. In this study, we chose PATI-treated leaves and floral organs as models of leaves with altered morphology and venation pattern; and investigated the spatial position of cell proliferative area, cell division angle, and possible factors that control the properties of lateral organ meristems using AN3 as a key clue.

## Results

### Change in the length of cell proliferation zone

Firstly, we examined whether PATI treatment affects leaf shape via changes in leaf meristem positioning in *A. thaliana.* In the PATI-treated plants, the leaves became shorter and rounder than those of the control plants (**Figs. 1D-F, 2**). The leaf vein pattern also differed in PATI-treated leaves (**Fig. 1D-F**); namely, the midvein was widened, the lateral veins ran parallel to each other, and the veins adjacent to leaf margins were also widened, indicating drastic changes in leaf organogenesis. These observations are consistent with those of previous reports (Mattsson et al., 1999; Sieburth, 1999; Sawchuk et al., 2013).

**Fig. 1.**
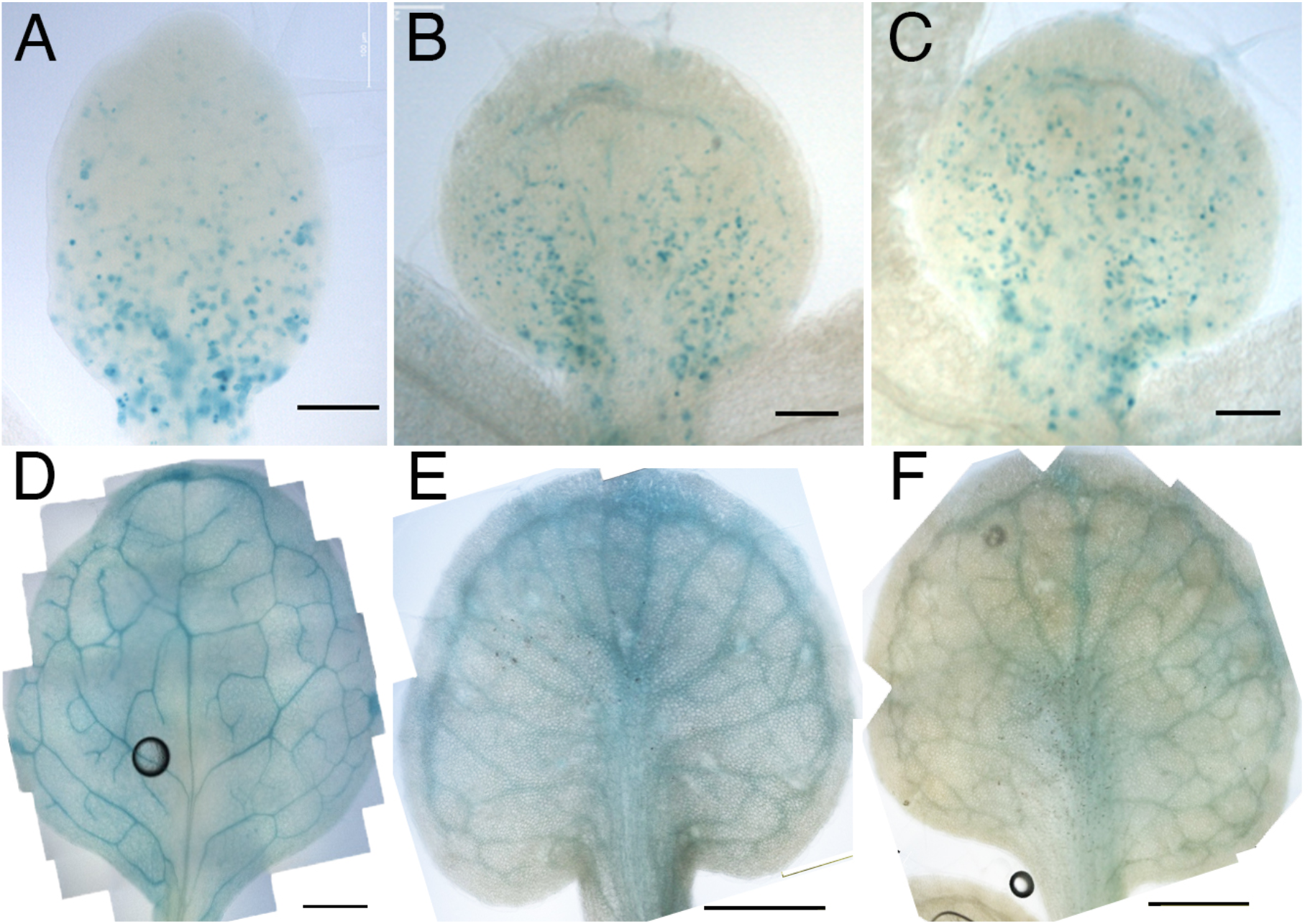
Cell proliferation is visualized by GUS staining in PATI-treated *CYCB1;1::GUS*. A: 6 DAS control, B: 7 DAS TIBA-treated, C: 7 DAS NPA-treated leaf primordia. D: 12 DAS control, E: 12 DAS TIBA-treated, F: 12 DAS NPA-treated leaf primordia. Leaf primordia of similar lengths were compared. Scale bars: A: 100 μm, B, C: 200 μm, D-F: 500 μm

**Fig. 2.**
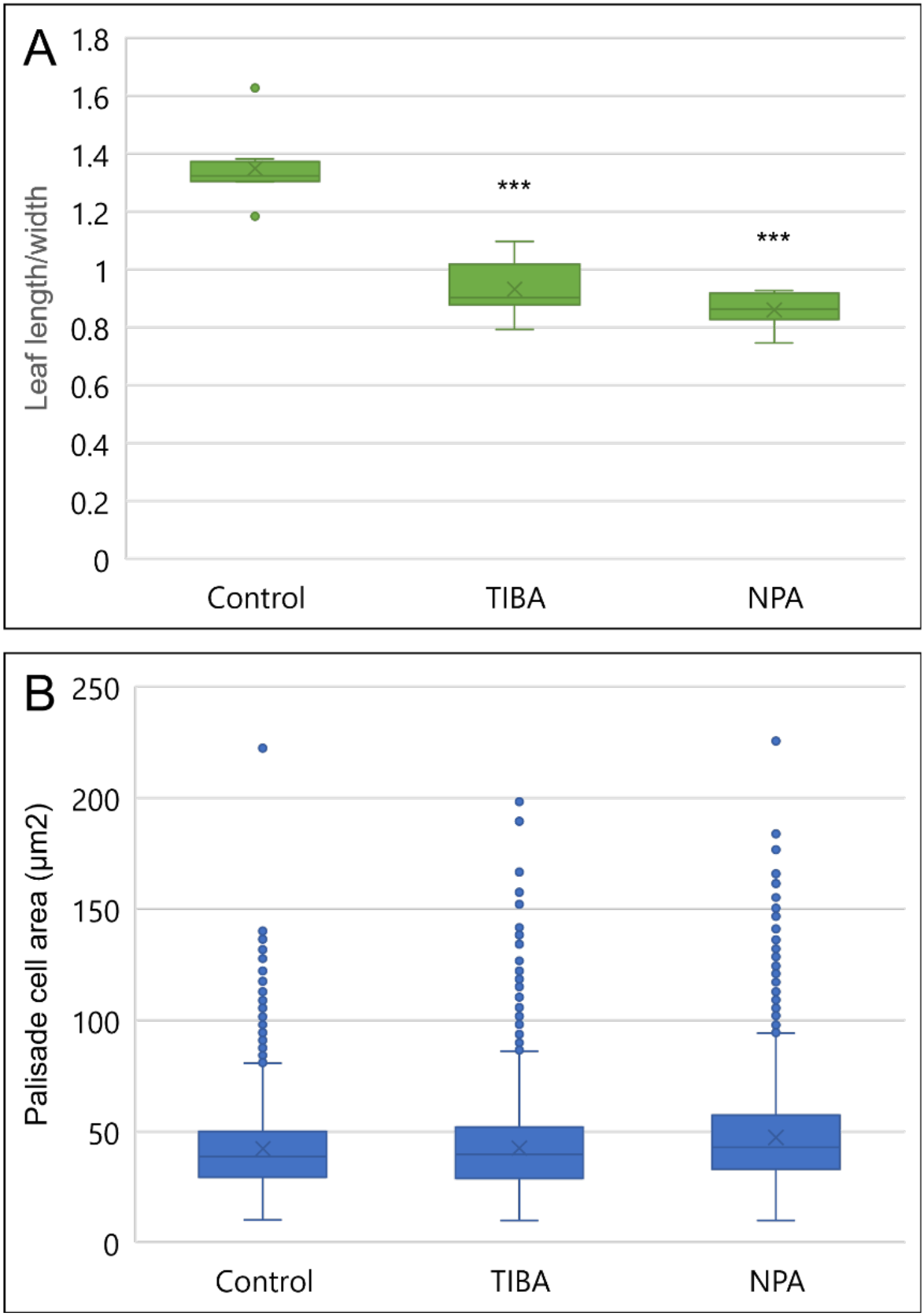
A: Leaf length/width ratio of leaf primordia at cell proliferation stage. Leaf length was measured after excluding the petiole, as shown in Figure 3A. Leaf width was measured after the leaf has been flattened out on glass slides. From left to right, data on control (6 DAS), TIBA-treated (7 DAS), and NPA-treated (7 DAS) leaf primordia are shown, n = 4, Dunnett’s test ***: *p* < 0.001 B: Palisade cell area for each condition. From left to right, data on control (6 DAS), TIBA-treated (7 DAS), and NPA-treated (7 DAS) n = 4

To investigate the effect of inhibition of PAT on the leaf meristem, the cell proliferation zone was observed in the leaf primordia of PATI-treated plants. The *CYCB1;1::β-glucuronidase* (*GUS*) line is used to visualize dividing G2-M phase cells (Donnelly et al., 1999). The first and second rosette leaves of 6 days after sowing stage (DAS) seedlings for control plants and 7 DAS seedlings for PATI-treated plants were used for this experiment, considering the growth retardation observed in the PATI-treated plants.

In the PATI-treated plants, the cell proliferation zone remained in the proximal position, similar to the control plants (**Fig. 1A-C**). The images were processed to further examine the positioning of the cell proliferation zone (**Fig. 3**). We observed that both the length of the cell proliferation zone from the leaf base and the ratio of cell proliferation zone to the total leaf length were found to be increased in PATI-treated leaves (**Fig. 3C, D**).

**Fig. 3.**
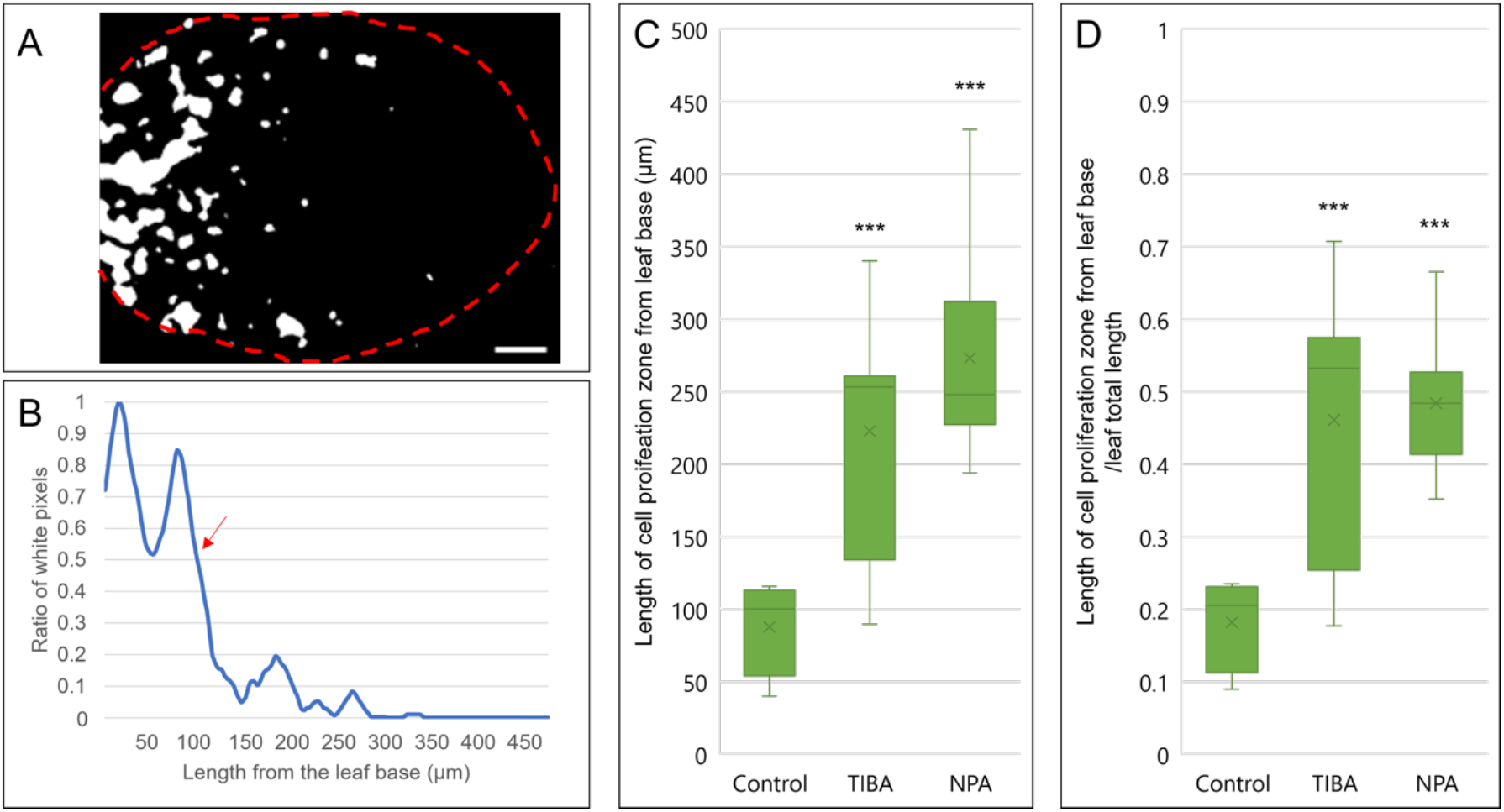
Determination of cell proliferation zone and length and the ratio of cell proliferation occupying leaf primordia. A: Binary images of GUS-stained leaf primordia. Scale bar: 50 μm. B: The ratio of white pixels from a binary image was calculated, and the length of the cell proliferation zone (more than 50% of the white pixels) was determined. The red arrow indicates the end of the cell proliferation zone. C: Length of cell proliferation zone from the leaf base. D: The ratio of cell proliferation zone to the total leaf length. n = 8, Dunnett’s test, ***: *p* < 0.001

### *AN3* mRNA and AN3 protein localization in leaf primordia

To investigate how the length of the cell proliferation zone increased, we examined the localization of *AN3* expression. As AN3 protein can move between cells, two lines, *an3-4*/*pAtAN3::AN3-GREEN FLUORESCNT PROTEIN* (*GFP*) and *an3-4*/*pAtAN3::AN3-3xGFP*, were used for the observation. The former can detect actual protein localization, whereas the latter is used to monitor mRNA localization because it represents an accumulation of the protein without cell-to-cell movement ability (Kawade et al., 2013, 2017). The first and second rosette leaves of 5 DAS seedlings were used for this analysis.

We observed that the overall localization of AN3 under PATI treatment did not change from that of the control; it remained in the proximal region (**Fig. 4A-L**). This confirms the observation in the cell proliferation zone described above. The size of the *AN3*-expressing regions was also measured using the same image processing method used for the analysis of the cell proliferation zone. Consequently, the *AN3*-mRNA expressing regions, monitored by the presence of the AN3-3xGFP signal, were slightly longer in the PATI-treated leaves than in control (**Fig. 4M**). However, AN3 protein localization of TIBA- or NPA-treated plants did not show a statistical difference from that of control plants (**Fig. 4M**). Therefore, irrespective of changes in *AN3* mRNA expression pattern, the spatial gradient of AN3 protein along the longitudinal axis of leaf primordia did not change. Therefore, the influence of PATI treatment on the AN3 protein accumulation pattern is rather limited.

**Fig. 4.**
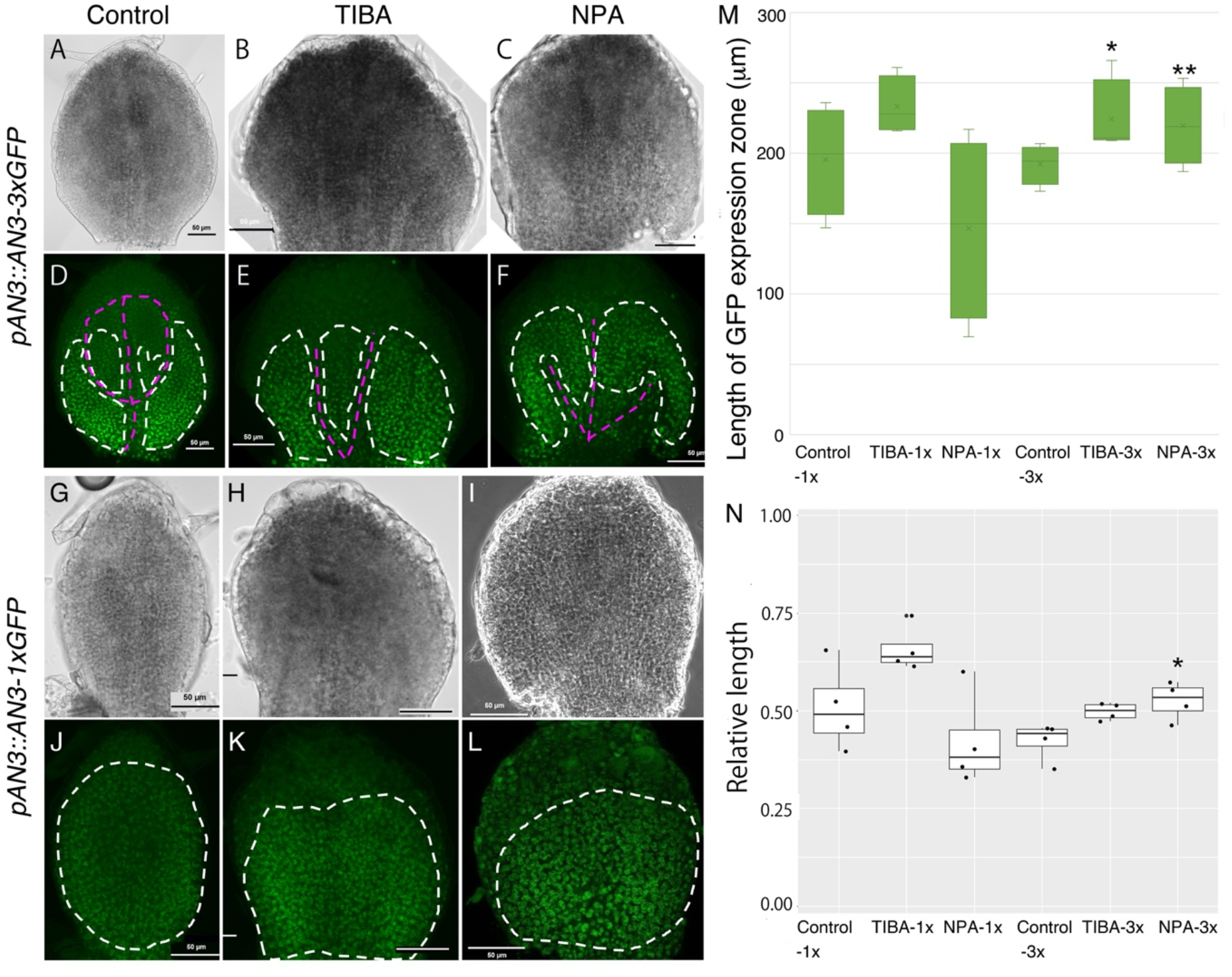
GFP localization of *an3/pAtAN3::AN3-3xGFP* (A-F) and *an3/pAtAN3::AN3-GFP* (G-L), and their ratio of AN3-3xGFP and AN3-1xGFP localized area to the total length of leaf primordia. Control (A, D, G, J), TIBA-treated (B, E, H, K), and NPA-treated (C, F, I, L) leaf primordia (5 DAS) are shown. White dotted lines indicate GFP-localized area, and magenta lines indicate vasculature. M indicates the actual length of GFP expression zone measured from leaf base. From left, *an3/pAtAN3::AN3-GFP* Control, TIBA and NPA; *an3/pAtAN3::AN3-3xGFP* Control, TIBA and NPA. n = 4, Dunnett’s test, *: *p* < 0.05, **: *p* < 0.001 N indicates a ratio of AN3 mRNA and protein-localized area to the total length of leaf primordia. From left, *an3/pAtAN3::AN3-GFP* Control, TIBA and NPA; *an3/pAtAN3::AN3-3xGFP* Control, TIBA and NPA. n = 4, Dunnett’s test, *: *p* < 0.05. No mark implies no statistically significant difference was observed.

On the other hand, when *an3-4*/*pAtAN3::AN3-3xGFP* were treated with PATI, we recognized that *AN3* localization was missing in the vasculature region. This missing localization was also observed in control conditions, but in PATI-treated leaves, the vasculature was very thick, and therefore, the absence was recognizable (**Figs. 1D-F, 4**).

### Cell division angles in leaves

To understand how PATI treatment caused the changes in size or pattern of the cell proliferation zone and the leaf shape, analysis of cell division angles was performed using the *gl1-s92f* mutant and *gl1-s92f*/*an3-4* double mutants. We chose *gl1-s92f* mutants that lacked trichomes as a control WT because trichomes obstruct cell division plane observation using aniline blue staining (**Fig. S1**). The angles were determined against the proximal-distal axis, starting from the leaf base and parallel to the midvein for the first and second rosette leaves of 7 DAS seedlings (**Fig. 5**). As a result, a variation in cell division angle patterns was observed (**Fig. 5A**). In the control WT plants, the cell division angle peaked at around 130°–140° (**Fig. 5A**). In PATI-treated leaves, this peak was less evident, and cell division was likely randomized (**Fig. 5A**). This may have contributed to changes in leaf shape, with rounder and shorter leaves, when treated with PATI (**Figs. 1D-F, 2A**).

**Fig. 5.**
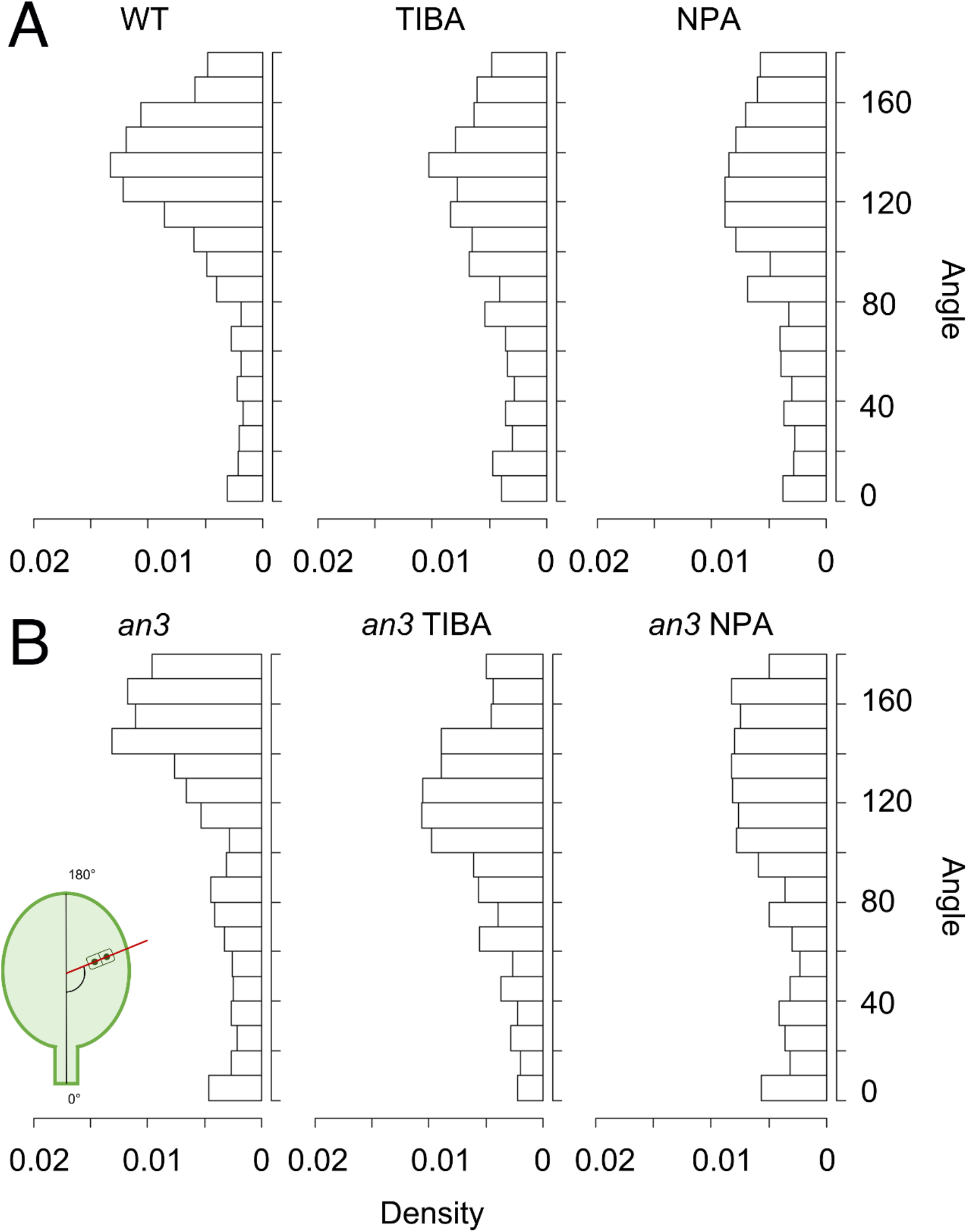
Cell division angles in leaves. Four samples were investigated for each condition. A: From left, WT, TIBA, NPA; 1221, 1158, 1061 pairs of cells were analyzed, respectively. B: From left, *an3* mutant, TIBA (*an3* mutant), NPA (*an3* mutant); 971, 703 and 604 pairs of cells were analyzed, respectively. A schematic view of the angle measurements is inserted at the lower left.

We performed a cell division angle investigation also for the *an3-4* mutant plants. In the *an3-4* mutant without PATI, the peak of cell division angle was around 140–150°, which differed from that of the WT (**Fig. 5A, B**), suggesting that AN3 might shift the cell division angle to smaller angles, i.e., a shift from proximo-distal to mediolateral direction. This angle shift can explain why the *an3* leaves are narrower than WT leaves. In the *an3* mutant treated with PATI, however, the distribution of cell division angle became similar to that of WT plants treated with PATI (**Fig. 5A, B**), indicating that the randomizing effect of PATI on cell division angle is superior to the effect of the *an3* mutation biasing cell division angle to the proximo-distal axis. In both WT and *an3*, the effects of PATI on the cell division angle seem to be similar between TIBA and NPA, but slightly stronger in NPA than TIBA (**Fig. 5A, B**).

### The position of cell proliferative area in floral organs

Then, we examined floral organs as modified leaves. Although floral organs such as sepals, petals, stamens, and carpels are homeotic to leaves, their shapes are different. Even between sepals and petals, both of which are planar organs, there are distinct differences in shape in *A. thaliana*. For example, in the distal part, the sepal is narrower, and the petal is wider, while foliage leaves are narrower in the distal part, similar to sepals.

We observed cell proliferation patterns in planar floral organ primordia to investigate whether these patterns might influence the difference in the final organ shape. To visualize dividing cells in floral organs, we used EdU. In a sepal primordium, cell division was observed in the basal part of the organ primordia through the observed developmental stages (**Fig. 6**). The position of cell proliferative area was similar to that of leaf primordium investigated in previous studies (e.g., Donnelly et al., 1999, Kawade et al., 2017) (**Fig. 1**). In contrast, cell division in the petal primordia was observed in the whole organ when the organ was around 100–150 μm in length and the distal and marginal regions when the organ was around 400 μm in length (**Fig. 6**), which marked a clear difference from that of leaves and sepals.

**Fig. 6.**
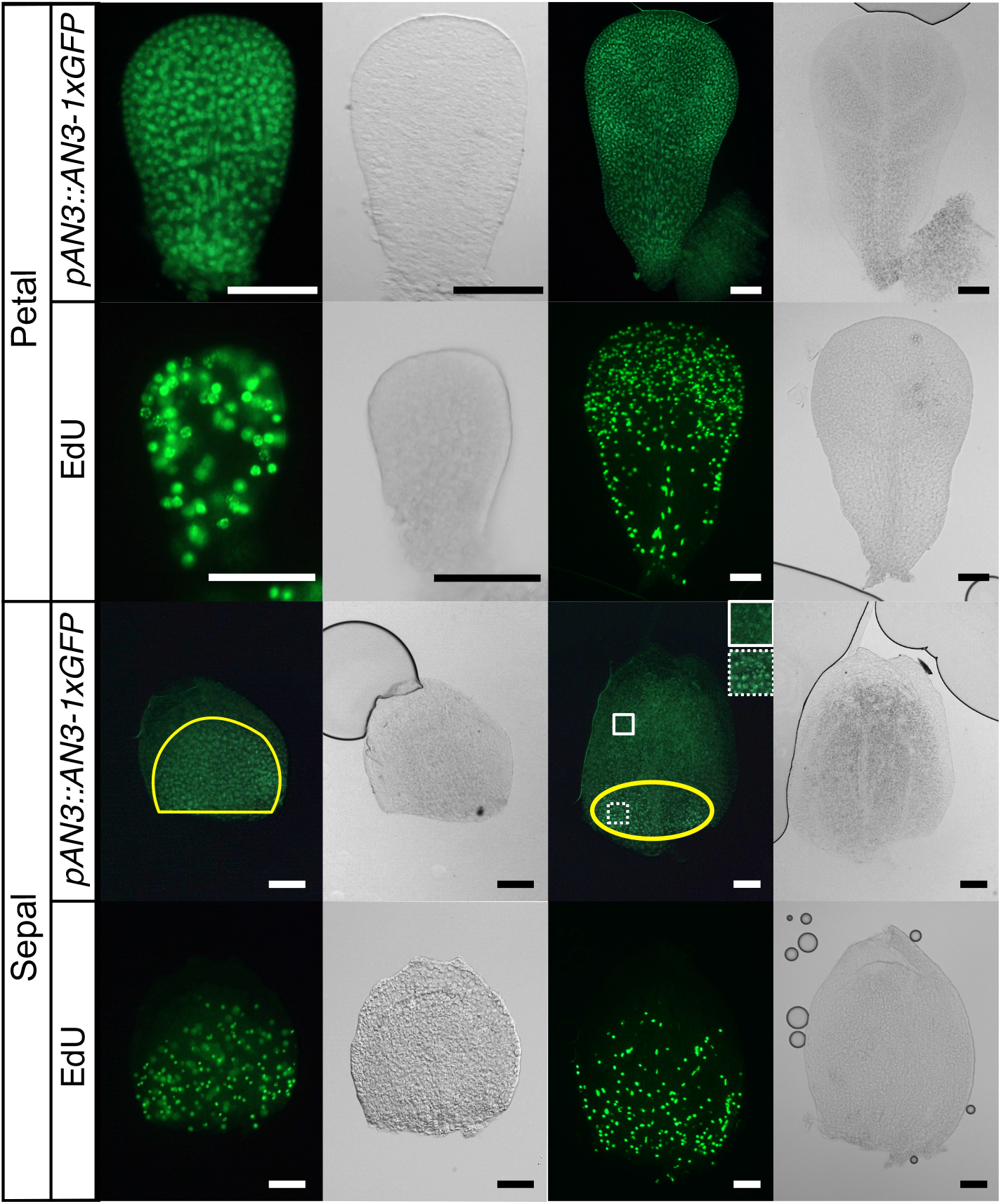
AN3-1xGFP signals and EdU staining in sepal and petal primordia. The primordia in the left two columns are younger than that in the right two columns. The stage of flower from which each primordium was collected is as follows: petal left, early stage 9; petal right, stage 1; sepal left, stage 7 (approx. 250 μm); sepal right stage 8 (approx. 450 μm). Normal transmitted light microscope images are shown on the right side of each fluorescent microscope image. Images of an AN3-GFP line are in the first row of each floral organ. The second row shows the floral primordia from the wild type treated with EdU. The yellow line shows areas with brighter AN3-1xGFP signals than the other part of the area in the primordia. Magnified views in each square of the sepal primordium are shown on the upper right. Bar = 50 μm

### AN3 protein localization in floral organs

As AN3 is a key regulator of the leaf meristem position, we suspected that AN3 protein accumulation patterns may be different between leaves and planar floral organs, which have different positions of the cell proliferative area. AN3-GFP signals were only observed in the basal part of sepal organ primordia, whereas the signals were observed in the entire petal organ primordia at first, and then in the distal region in later stages. Moreover, in the petal primordia, sparse signals were also observed in the central region and the basal part in the later stage 9 (**Fig. 6**), where EdU signals were rarely observed.

### Phenotype of *an3* mutant in sepal

It is reported that *an3* mutants have narrower petals and a smaller number of cells than that of the Col-0 wild type (WT) (Horiguchi et al., 2005; Lee et al., 2009), but the sepal phenotype has not been investigated. The above-mentioned similarity in the AN3 protein accumulation pattern and proliferative area in the sepal primordia strongly suggested that the *AN3* is also involved in meristematic regulation in the sepal. In order to investigate whether AN3 is involved in meristematic regulation as in leaf meristem in sepal primordia, we compared the phenotype of *an3-4* and the WT in the sepals (**Fig. 7A, B**). The area of the organ was smaller in *an3* than in the WT (**Fig. 7C**). We also observed that the *an3* mutant had fewer complex veins as compared to that of the WT, which was evaluated based on the number of secondary and higher veins in the m-shaped or two n-shaped primary veins (**Fig. 7D, E**).

**Fig. 7.**
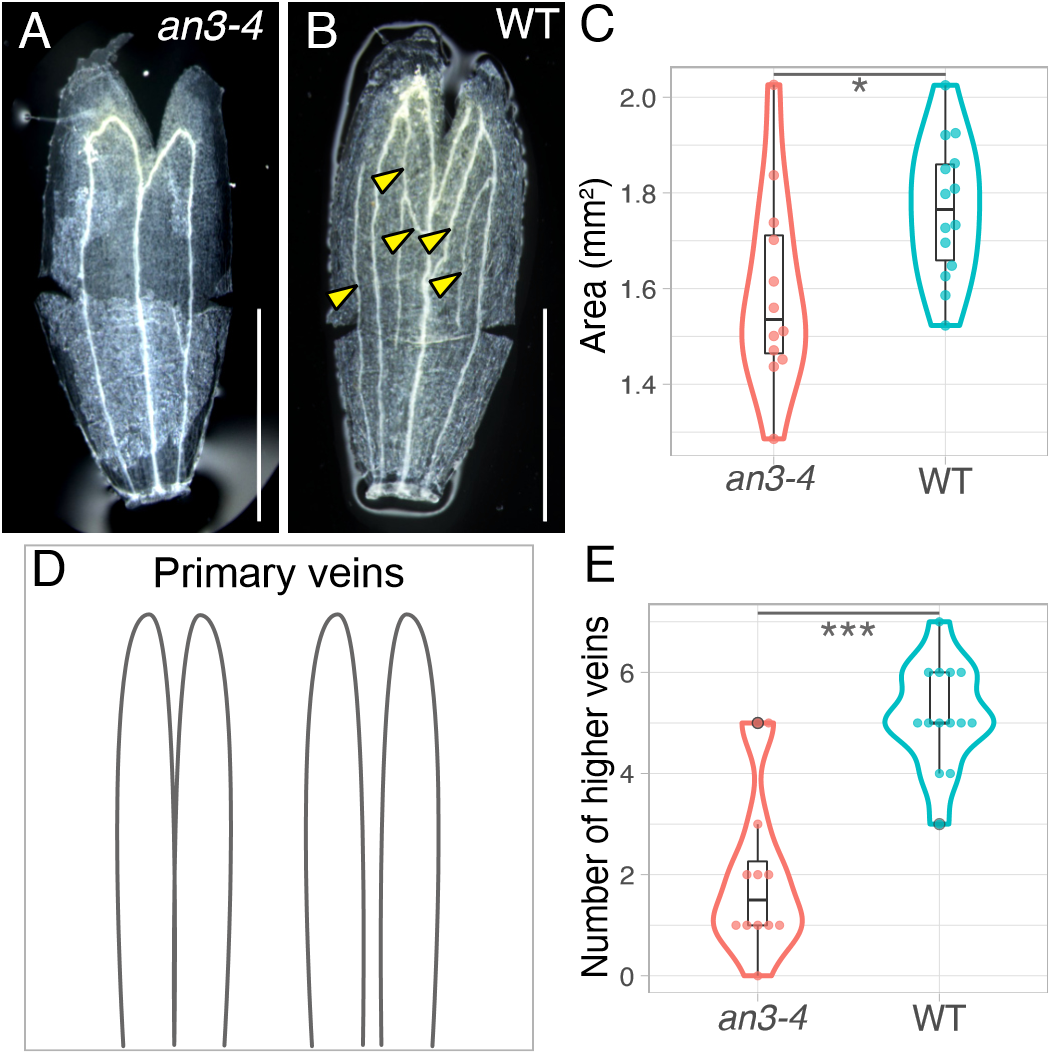
Phenotype of the sepal in the *an3-4* and the wild type. (A, B) Images of the sepal in each genotype, *an3-4*, and the wild type. Some cuts were made to flatten the organs. The yellow arrowheads point to the higher veins in the sepal. (C) The area of sepals in each genotype. (D) Two patterns of primary veins are defined in this study. (E) The number of higher veins of the sepals in each genotype. * *p* < 0.05, *** *p* < 0.001, Bar = 500 μm.

### Cell division angles in floral organs

To know the possible contribution of biased cell division angle on the final shape of floral organs, cell division angle analysis was conducted in the sepals and petals from flowers at stages 8–10 where active cell proliferation is known (Alvarez-Buylla et al., 2010). The distribution pattern of cell division angle in planar floral organs was unique in showing loose double peaks (**Fig. 8A**) that is different from the cell division angle in leaves, which had one peak (**Fig. 5A, B**). Irrespective of the similarity between sepals and leaves in terms of localization of the cell proliferation zone, the overall tendency of cell division in sepals was similar to that of petals. This suggests that the pattern of cell division angles is not associated with the localization of the cell proliferation zone but with organ identity.

**Fig. 8.**
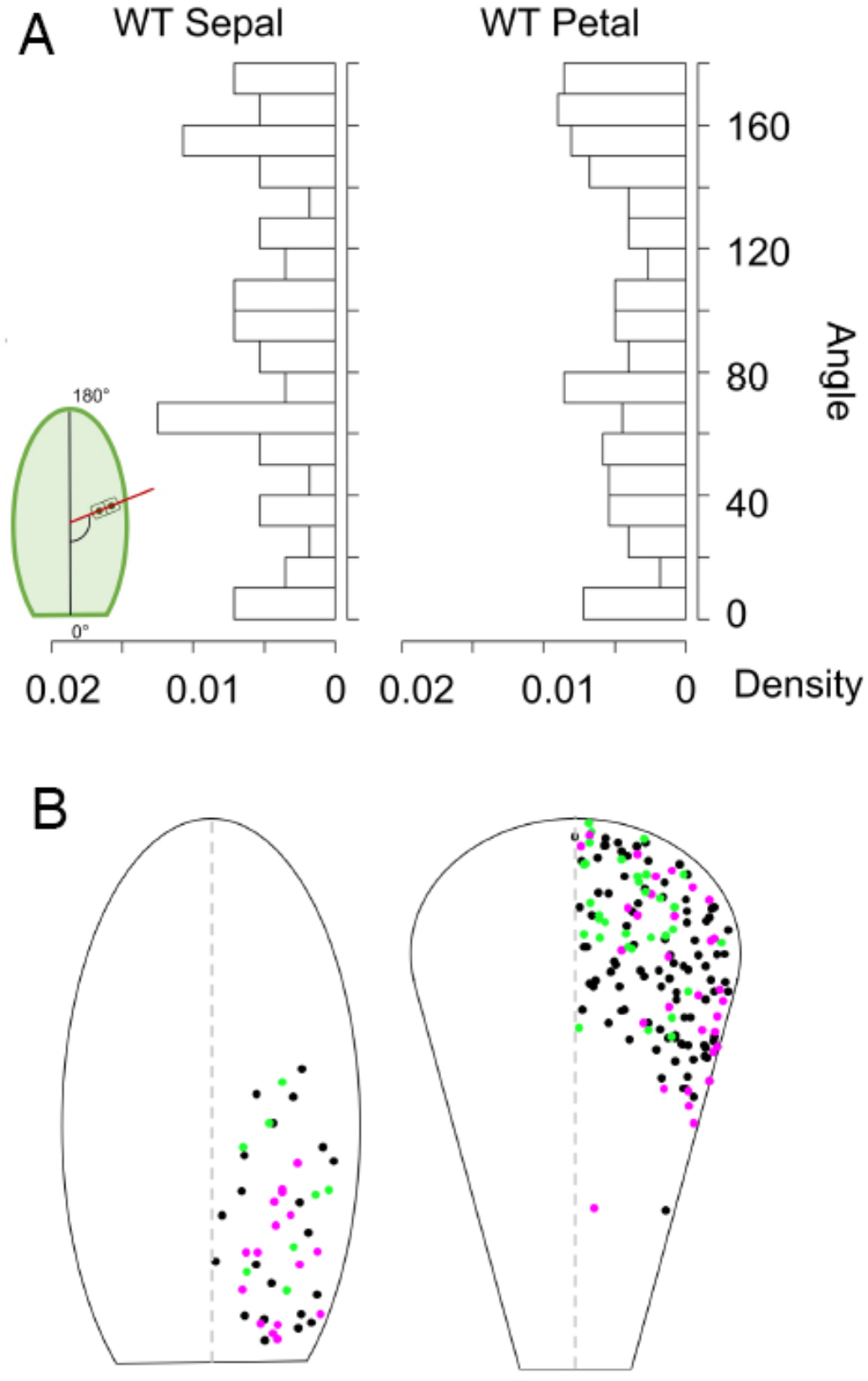
Cell division angles in floral organs. (A) From left, WT sepal, WT petal; 56 and 222 pairs of cells from four samples were analyzed, respectively. A schematic view of the angle measurements is inserted in the left corner. (B) Cell division angle distribution in floral organs. Left: sepal; 46 pairs of cells from three primordia were analyzed. Right: petal; 159 pairs of cells from three primordia were analyzed. The angles are mapped to the right half of the diagram, as the organs are symmetrical. The magenta dots indicate the cell division angle in 140°–180° (the upper peak in the panel A), the green dots indicate 60°-90° (the lower peak in the panel A), and the black dots indicate the angles that are in neither peak. n = 3

To understand further the unique cell division patterns in these floral organs, we also analyzed the spatial distribution of the cell division angles in the primordia (**Fig. 8B**). As a result of the sepal primordia, the cell divisions within the 60°–90° range, which were divisions in the mediolateral direction of the primordia, were identified more distally than the divisions that corresponded to the 140°–180° range, which were divisions in the proximo-distal direction (**Fig. 8B**), which may contribute to the oval-shaped sepal primordia. On the other hand, in the petals, the cell divisions within the 60°–90° range, which were divisions in the mediolateral direction of the primordia, were identified mostly in the central regions of the petal primordia. The divisions that corresponded to the 140°–180° range were identified mostly in the marginal regions of the petal primordia (**Fig. 8B**).

## Discussion

In this study, we examined how cell proliferative zones differ in the primordia of PATI-treated leaves and floral organs from normal foliage leaf primordia of *A. thaliana* with a focus on the spatial expression pattern of *AN3*, a key element for leaf meristem positioning (Kawade et al., 2010). We identified that organ shape variation by PATI treatment cannot be attributed to changes in leaf-meristem positioning, size of the leaf meristem, or the expression pattern of *AN3* but is rather attributed to altered cell division angles in the leaf meristem. Interestingly, the *an3* mutation biased cell division angle to the proximo-distal axis in the leaf meristem, whereas PATI treatment randomized the angle. These effects on the cell division angle are reasonable considering that the *an3* has narrower leaves and PATI-treated leaves are shorter and rounder (**Figs. 1, 2, 4**). Because polar auxin flow is highly plausible to polarize the cell division angle (Yin and Tsukaya 2016; reviewed in Dhonukshe et al. 2005), the observed phenomena could be explained as follows: AN3 randomized the cell division angle (or shift the angle from proximo-distal to mediolateral direction) against the auxin-dependent polarity; PATI randomized the cell division angle by cancelling the auxin flow. Since along the vasculatures *AN3* mRNA expression was not observed (**Fig. 1D-F**), polarized cell division angle along the vasculature in the WT (Yin and Tsukaya 2016) could be explained by the absence of AN3.

Different shapes of sepals and petals compared with foliage leaves were found to be correlated with both altered meristem position associated with altered *AN3* spatial expression patterns and different distributions of cell division angles. Overall, our results strongly suggest that lateral organ shapes are regulated via two aspects: position of meristem and cell division angles; the former is mainly governed by the *AN3* expression pattern. In the following sections, several aspects of the above findings are discussed.

### The position of leaf meristem in PATI-treated plants

When *A. thaliana* plants were treated with PATI, both the cell proliferation zone and *AN3-*expression zone still sit in the proximal par of leaf primordia. It was also observed that under the PATI treatment the *AN3* mRNA expression zone was slightly expanded to the distal direction in the expression zone ratio, although the final AN3 protein distribution remained unchanged (**Fig. 4M, N**). As previously reported (Sieburth, 1999) and confirmed here (**Figs. 1-3**), PATI-treated leaves were rounder and shorter, whereas a longer proliferation zone would be expected to produce a longer leaf if the other aspects were not changed. Instead, we found that changes in the cell division angles could be attributed to the altered leaf shape.

We also observed that the *AN3* promoter was not expressed in leaf veins (**Fig. 4A-L**). This trend was clearly seen in PATI-treated leaves as well as in control conditions; however, it was not evident in control conditions because the veins were much thinner than those of PATI-treated leaves (**Fig. 1D-F**). This may happen if *AN3* expression is shut down in differentiated vascular cells, which may imply that the vasculature differentiation by auxin is superior to cell proliferation maintenance by AN3.

### Cell division angle and leaf shape

Cell division is an important factor in both leaf development and leaf vein architecture (Kang et al., 2007). In this study, analysis of the cell division angle revealed that the pattern differed between PATI-treated plants and control plants (**Fig. 5A**). This difference in the pattern could be a cause for the differences in leaf shape. In comparison with control leaf primordia that had major division angles in 130°–140°, PATI-treated leaf primordia had dispersed division angles in many directions, forming a round and short leaf, which matched the phenotype (**Figs. 1D-F, 3**). This suggests that auxin flow regulates leaf shape via controlling cell division orientation in the leaf meristem. It has been shown that the presence of vasculatures is important in determining cell division patterns (Yin and Tsukaya, 2016). Therefore, the thickened midveins, where many veins running in the same direction along proximal-distal axis observed in PATI-treated leaves, may have indirectly affected the cell division angle.

### Determining cell division angle in leaves

In this study, two components changed the cell division angle: PAT and AN3. When PAT was inhibited, the peak in the cell division angle distribution became less evident (**Fig. 5A**). As auxin flow controls vascular cell polarity (Linh et al., 2018) it is possible that the cells divide in the direction of auxin flow. In addition, both NPA and TIBA affect actin dynamics (Teale and Palme, 2018; Zou et al., 2019) that can affect cytoskeletal regulation of cytokinesis, and may be one of the underlying reasons for the change in cell division angle.

We observed the *an3* mutant tended to divide around 140°–150°, which partially explains the narrow leaf phenotype of *an3* mutants, as cell division along the proximal-distal axis was increased (**Fig. 5B**). In a previous report, the phase of cell division was divided into two phases: the first has more divisions along the proximal-distal axis than the second phase (Horiguchi et al., 2011). In the latter phase, except for marginal area and local areas along veins cell division orientation was randomized (Yin and Tsukaya 2016). Furthermore, in *an3* mutants, the transition from the first to the second phase does not occur before the termination of cell division activity (Horiguchi et al., 2011). The shift in the peak of the cell division angle observed in the *an3* mutant would reflect this failure in shifting to the second phase of cell proliferation, which confirms the results of previous studies. Therefore, AN3 functions in the transition to the second phase of cell division, and consequently, cells tend to divide along the proximal-distal axis in the absence of AN3. Precisely, AN3 might promote the shift of the cell division angle from the proximo-distal preference to the randomized one.

In addition, in *an3* mutants treated with PATI, the cell division angles were similar to those of WT treated with PATI (**Fig. 5A, B**). This suggests that the randomizing effect of PATI on cell division angle is superior to that of AN3 and that the loss of PAT results in cell division in random directions irrespective of the presence or absence of AN3. This might be explained as follows: AN3 functions for the phase shift from proximo-distal preference to randomized one; polar auxin transport is directly or indirectly involved in the polar-dependent cell division angle in both phases; loss of AN3 activity and PAT results in more proximo-distal and randomized cell division, respectively. If the PAT-dependent polarity axis is lost, even in the *an3*, the cell division angle is randomized.

### Position of meristematic tissue determines the final floral organ shape

The final organ shape could be determined by various factors, such as acceleration and deceleration of proliferation, oriented cell division and expansion, and the meristem position (Tsukaya, 2018). In this study, we showed that the position of the cell proliferative area was completely different between sepals and petals, which are both modified leaves but morphologically different. The petal of *A. thaliana* with a modest fan shape has a proliferative region in the distal part, which is similar to ferns with fan-shaped morphology, which is rare in angiosperm leaves, coinciding with leaf meristem at the apical margin (Boyce, 2007; Tsukaya, 2014, 2018). This suggests that the morphological differences between sepals and petals could be at least partly explained by the meristematic position in each organ.

Although predominant cell division occurs in submarginal plate meristem to widen the leaf blade area in leaf primordia (e.g., Poethig and Sussex, 1985), it was also shown that marginal meristem residing in the margin of the primordia is present (Alvarez et al., 2016; reviewed in Tsukaya 2021). Alvarez et al. (2016) showed that when NGATHA and CINCINNATA-class-TCP were knocked down, indeterminate marginal growth occurred in the entire margin of the leaf blade, suggesting potential meristem activity in this area. Interestingly, marginal growth occurs only in the distal region of floral organs in their system, including sepals and petals. In terms of WT petals, active cell division occurs in the distal margin in the first place (Sauret-Güeto et al. 2013; our present study, **Fig. 6**); therefore, the marginal meristem may contribute more than leaves or sepals. Since the activity of meristem in the distal marginal area of a blade has been discussed as an ancestral character (Floyd and Bowman, 2010), petals may retain this developmental character (Boyce, 2007). We observed EdU signals in the entire margin of the petal primordia at least until the organ size was 400 μm in length, even in the proximal region. Considering the results of Alvarez et al. (2016) on leaf and floral organ primordia, the nature of proliferative cells in marginal areas is different among different lateral organs.

### Role of AN3 in planar lateral organs

Several key genes are known to positively control cell proliferation in the leaf meristem (Nakata et al., 2012; Ichihashi and Tsukaya, 2015; Tsukaya, 2021). Among them, the AN3-protein-accumulated region matches with the cell proliferative area in leaf primordia (Kawade et al., 2017), suggesting that AN3 is an important determinant of the leaf meristem position. Moreover, the smaller size of a petal in *an3* or triple knockdown of the GIF family (Kim and Kende, 2004; Horiguchi et al., 2005; Lee et al., 2009) suggested that AN3 is also involved in the promotion of cell proliferation in the petal. However, its functioning zone in primordia has not been well investigated. In this study, we showed that the AN3-expressed region overlapped with the cell proliferative area in both sepals and petals, as observed in leaf primordia. In addition, we first showed that traits of sepals in *an3* mutants are likely to have fewer cells as petals or foliage leaves. These results suggest that AN3 functions as a determinant of the meristematic position and activity in all planar floral organs. However, in terms of petal primordia, AN3 protein signals were also observed in the less proliferative area. This might be due to a lack of associating transcriptional factors such as GROWTH REGULATING FACTOR5 (GRF5), which is necessary to promote cell proliferation in leaf primordia (Horiguchi et al., 2005), in such regions. Alternatively, because signals in the proximal part were not as strong as those in the distal region, the concentration of AN3 proteins might not be enough to promote cell division to the extent that EdU was incorporated.

In the past, JAGGED (JAG) was examined as a candidate of a regulatory gene for the specific morphology in the *Arabidopsis* petal that differed from leaves because the loss-of-function *jag* mutant has narrower and shorter petals; the *JAG* overexpressor has larger petals; its mRNA is expressed distal margin (Sauret-Güeto et al. 2013). At that time point, JAG was the only candidate ‘organizer’ that presented the petal with a pattern of growth orientations that fans out. However, AN3 has become an additional candidate that fulfills the required, above-mentioned conditions. Indeed, *AN3* was identified as a direct target of JAG (Schiessl et al. 2014). The role of AN3 as the ‘organizer’ sensu Sauret-Güeto et al. (2013) should be examined in the future.

To fully understand the mechanisms of flower organogenesis, the regulation of meristem position by floral organ identity genes needs to be investigated (Coen and Meyerowitz, 1991). Honma and Goto (2001) and Pelaz et al. (2001) revealed that when A genes (*APELATA1*) and B genes (*APELATA3* and *PISTILATA*) were ectopically expressed together with *SEPALLATA2/3* in rosette leaves, the rosette leaves obtained petal identity, and the color and cell shape became petal-like. However, the transformed petaloid organ was not fan-shaped but had a taper off shape, which was similar to rosette leaves, cauline leaves, and sepals. This suggests that factors other than the floral identity homeotic genes control the final organ shape. Revealing the mechanisms of *AN3* expression control might shed light on which factors are involved in the resulted different morphology among different organs.

The leaves of some gymnosperms and ferns are considered to grow from the meristem in the distal margin. The positioning of these meristems may also be determined by the spatial distribution of leaf meristem-controlling genes, such as *AN3/GIF1*. As *GIF* family genes exist in most eukaryotic organisms, including the basal land plants, *Marchantia polymoprha*, *Physcomitrium patens*, and *Sellaginella moellendorffi* (Kim and Tsukaya, 2015), further analyses of the *GIFs* in gymnosperms and ferns could answer this question.

### Determining cell division angle in floral organs

In this study, the cell division pattern in floral organ primordia was investigated for its possible roles in each floral organ morphology. Both the petals and sepals showed a pattern with loose twin peaks in the distribution of cell division angles, which was different from that of leaf primordia that had a clear single peak (**Fig. 6**). This is an interesting new finding on the meristems in these primordia. In addition, we found that in the petal the cell division occurred at 60°–90° angle in the central regions, and the cell division with the 140°–180° angle was mostly in the marginal regions, whereas such a pattern was not seen in the sepals (instead longitudinal distribution was observed, **Fig. 8**). This difference may cause variation in shape between sepals and petals, with the distal part being wider than the proximal part, which may be caused by the cell divisions that contributed to width in the marginal regions. Although both sepal and leaf primordia have a cell proliferation zone in the basal region, the cell division angle was controlled differently, which may imply that cell division angles depend on organ identity and affect their final shapes.

Overall, our results imply that lateral organ shapes are likely to be regulated by two factors: the position of the cell proliferative zone governed by the spatial expression pattern of *AN3* and cell division angles. Therefore, even in one species, by changing these two factors, a variety of lateral organ shapes could be performed. To test this idea, future studies with a combination of genetic manipulation and computer simulation should be carried out.

## Materials and Methods

### Plant growth

For analysis of the leaf primordia, *A. thaliana* Col-0 (WT), or those carrying CYCLINB1;1(CYCB1;1)::GUS, an3-4, an3-4/pAtAN3::AN3-GFP, an3-4/pAtAN3::AN3-*3xGFP*, *gl-s92f*, or *gl-s92f*/*an3* were grown on sterile growth medium that contained half-strength Murashige and Skoog medium (MS, Wako, Osaka, Japan), 1% (w/v) sucrose (Nacalai Tesque, Kyoto, Japan), and 0.8% (w/v) agar (Nacalai Tesque, Kyoto, Japan) (Wako, Osaka, Japan) adjusted to pH 5.8 with potassium hydroxide. Approximately, 1 M PATI (TIBA and NPA) stocks were dissolved in dimethyl sulfoxide and added to the medium to a final concentration of 10 μM. The medium composition was based on that described by Sieburth (1999).

Seeds were sterilized by immersing in a solution of 2% (v/v) Plant Preservative Mixture™ (Plant Cell Technology, Washington, D.C., USA), 0.1% (v/v) Triton X-100 (Nacalai Tesque, Kyoto, Japan), and 50 mg/L MgSO_4_ for 6 h or a solution of 10% (v/v) sodium hypochlorite (Nacalai Tesque, Kyoto, Japan) and 1% (v/v) Triton X-100 for 5 min and twice with sterile water prior to plating. The plates were incubated at 24°C under constant illumination.

For analyses of the floral organ primordia, *A. thaliana* Col-0 and *an3-4*/*pAtAN3::AN3-1xGFP* (Kawade et al., 2013) were sown on rockwool (Toyobo, Osaka, Japan) and grown under white fluorescent light conditions (ca. 40 μmol m^−2^ s^−1^) at 22–23°C supplied with water containing 1 g/L powder Hyponex (Hyponex, Osaka, Japan).

### GUS experiments

Detection of GUS activity was carried out using 5-bromo-4-chloro-3-indolyl-β-D-glucuronide (X-Gluc) as a substrate. Plant tissue was first placed in 90% (v/v) acetone on ice for 10 min, washed with sodium phosphate buffer (pH 7.0), and then placed in X-Gluc buffer solution (0.5 mg/mL X-Gluc, 100 mM NaPO_4_ (pH 7.0), 5 mM K_3_Fe(CN)_6_, 5 mM K_4_Fe(CN)_6_, 10 mM EDTA, 0.1% (v/v) Triton X-100) under vacuum for 15 min or more and then placed in the dark at room temperature (ca. 20°C).

After GUS detection, plant tissues were rinsed in 70% (v/v) ethanol and fixed in ethanol: acetic acid (6:1) solution. After chlorophyll was removed, the tissue was preserved in 70% EtOH in the dark. Plant tissues were mounted on slides with chloral hydrate solution (50 g chloral hydrate, 5 g glycerol, 12.5 mL distilled water) (Tsuge et al., 1996) and observed under a microscope after the tissue became transparent enough.

### AN3-GFP observations

Leaf primordia (5 DAS) and flower primordia of *A. thaliana an3-4*/*pAN3::AN3-GFP* and *an3-4*/*pAN3::AN3-3xGFP* lines were fixed in 4% (v/v) paraformaldehyde (PFA) in phosphate-buffered saline (PBS) with 0.05% (v/v) Triton X-100 by immersing them in the fixation mixture, deairing them for 10 min (for leaf primordia) or 15 min (twice, for flower primordia) and placed at 4°C overnight. The samples were then washed in PBS (10 min, twice) and stored in PBS at room temperature for leaf primordia and 4°C for flower primordia until observation.

The samples were then dissected using a sharp razor under the microscope. Leaf primordia were mounted on slides with PBS and observed under a confocal microscope (FV3000; Olympus, Tokyo, Japan) with a GFP filter for leaf primordia and an upright fluorescent microscope (DM4000; Leica Microsystems GmbH, Wetzlar, Germany) for floral organ primordia.

### Data analysis on the area of cell proliferative area and AN3-GFP localized area

We used a method derived from Kazama et al. (2010) and Ikeuchi et al. (2011) to determine the position of leaf meristem. First, an image of a leaf with a GUS expression pattern was prepared. The outer region of the leaf was cropped, and the image was rotated so that the leaf base was on the left side of the image. Then, the blue region was extracted, and a binary image was created using ImageJ (https://imagej.nih.gov/ij/). The number of white pixels was counted for each column, and the end of the cell proliferative area along the proximal-distal axis (hereafter referred to as arrest front) was determined based on the definition of the point at which the ratio of white pixels was half that of the maximum and farthest from the blade base. The distance from the leaf base of each arrest front point was plotted for each condition in a box plot. Statistical analysis was performed using the R software.

Similarly, AN3-GFP localized area was determined as follows: first, a series of z-stack images were stacked using ImageJ software. Stacked images with the outer side of the leaf were cropped and rotated so that the leaf base was on the left side of the image. The region with AN3-GFP fluorescence was extracted, and from this image, a binary image was created. The number of black pixels was counted in each column. The end of the AN3-GFP localized area was determined based on the definition of the point at which the ratio of black pixels was half that of the maximum and farthest from the blade base. The distance from the leaf base of each end of the AN3-GFP localized area was divided by the leaf length because of the size difference between lines. The obtained data were plotted for each condition in a box plot. Statistical analysis was performed using the R software.

### Observation of Aniline Blue signal

We used a method derived from previous studies for the detection of newly formed cell walls (Kuwabara and Nagata, 2006; Kuwabara et al., 2011). Leaf primordia (7 DAS) of *A. thaliana glabra1(gl1)-s92f* and *gl1-s92f*/*an3-4* mutants and Col-0 flower petals and sepals were first fixed in a mixture of ethanol and acetic acid (4:1, v/v) for 30 min and then rinsed in 100% ethanol. Then, the samples were immersed in a mixture of ethanol and 100 mM phosphate buffer (pH 9.0; 1:1, v/v) for 30 min and then in 100 mM phosphate buffer (pH 9.0) for 10 min. Finally, the samples were immersed in a 0.02% (w/v) solution of aniline blue in 100 mM phosphate buffer (pH 9.0) for at least 7 days and up to 30 days at 4°C. Leaf primordia were mounted on slides with the staining solution and observed under a confocal microscope (FV10C-PSU; Olympus, Tokyo, Japan) under UV excitation with a DAPI (4’,6-diamidino-2-phenylindole) filter. The data were analyzed by taking each angle of the septum wall. Calculations were performed using Microsoft® Excel.

### Observation of EdU-marked cells

We used the Click-iT EdU Alexa Fluor 488 Imaging Kit (Thermo Fischer Scientific, Waltham, MA, USA) to visualize the cells in S phase. We dissected the inflorescence of *A. thaliana* into several pieces and soaked the flower clusters into 10 μM 5-ethynyl-2’-deoxyuridine (EdU) solution in half-strength MS medium with 1% sucrose for 3 h under ~45 μmol m^−2^ s^−1^ white fluorescent light. The samples were washed with PBS containing 0.1% Triton X-100 and fixed with FAA (37% [v/v] formaldehyde, 5% [v/v] acetic acid, 50% [v/v] ethanol) and stored at 4°C. Subsequent fluorescent labeling with Alexa Fluor 488 or 555 (Thermo Fischer Scientific, Waltman, MA, USA) was conducted according to the manufacturer’s instructions. Floral organs were mounted on slides, and fluorescent dye signals conjugated to EdU were observed under fluorescent microscope (DM4000; Leica Microsystems GmbH, Wetzlar, Germany).

## Acknowledgments

We would like to thank MEXT and the Graduate Program for Leaders in Life Innovation (GPLLI)/World-leading Innovative Graduate Study Program for Life Science and Technology (WINGS-LST) of the University of Tokyo for providing microscope facilities.

## Author Contributions

AK, MN, and HT designed the experiments; AK and MN performed the experiments and analyzed the data; AK, MN, and HT wrote the manuscript.

## Funding

This research was supported by a Grant-in-Aid for JSPS Fellows (AK, #19J14140) and a Grant-in-Aid for Scientific Research on Innovation Areas (HT, #25113002 and 19H05672) from MEXT and GPLLI/WINGS-LST of the University of Tokyo (AK).

## Conflict of Interest

No conflict of interest.

## Data Availability

Data is available on request from the authors.

## Notes

### Competing Interest Statement

The authors have declared no competing interest.

### Summary of Updates

Here some data were updated; figures were rearranged and revised; text was thoroughly rewritten; new references were added, too.

